# A fast and simple algorithm for accurate spike detection in HD-MEA recordings

**DOI:** 10.64898/2026.01.16.699955

**Authors:** Juan Zegers-Delgado, Nathaniel Renegar, Kasun Pathirage, Timothy K Horiuchi, Pamela Abshire, Ricardo C. Araneda

**Affiliations:** Department of Biology, University of Maryland, College Park, Maryland 20742, United States of America; Neuroscience and Cognitive Sciences Program, University of Maryland, College Park, MD, United States; Fischell Institute for Biomedical Devices, University of Maryland, College Park, MD, United States; Department of Electrical and Computer Engineering, University of Maryland, College Park, MD, United States; Institute for Systems Research, University of Maryland, College Park, Maryland, United States of America

**Keywords:** Spike Detection, Spike sorting, Multi-electrode array, High frequency firing

## Abstract

**Background:** High density microelectrode arrays provide a strong platform to study individual neuronal activity and neuronal network dynamics. However, the analysis of high volume and complex data present several challenges. Common spike detection methods based on Root-Mean-Square (RMS) threshold crossing underestimate the number of spikes during neuronal bursting, which frequently occurs in neuronal cultures. In addition, the detection of action potentials by multiple electrodes makes spikes sorting a computationally expensive task.

**New Method:** We optimized a previously described detection method, based on the scaled median of absolute deviations (MED) that is more accurate during high rates of neuronal firing. In addition, we added a step to de-duplicate (DP) spikes recorded on multiple electrodes, which enhanced the accuracy of MED. The combined method of detection and de-duplication (DP-MED) is less computationally expensive and easier to implement than popular sorting alternatives like Kilosort-4.

**Results and Conclusions:** During burst periods, the MED-based method detected over half of spikes that were undetected by the RMS-based method. To evaluate the performance of DP-MED, we simulated data that emulates neuronal activity recorded with HD-MEA. Across increasing firing rates, DP-MED shows more precision than Kilosort-4 but is slightly less accurate. After inducing high firing rate in cortical cultures with pharmacological stimulation, DP-MED detected a similar number of spikes than Kilosort-4, however, the analysis in Kilosort-4 was 40-fold more time-consuming. These results highlight the effectiveness of the DP-MED method in the context of drug screening using HD-MEAs.

## 1. Introduction

The combination of *in vitro* neuronal cultures with recordings in HD-MEAs is a powerful approach to study the basis for network dynamics in healthy neurons, aberrant neural function in disease models, and the discovery of potential treatments through drug screening (Hoang et al., 2025; Kobayashi et al., 2023; Duru et al., 2023). HD-MEAs consist of densely packed electrodes that allow the tracking of action potentials across the cell body of multiple cells with unsurpassed spatial resolution (Müller et al. 2015). Furthermore, several HD-MEA-based platforms allow long-term monitoring of neuronal activity, up to 40 days *in vitro* (Ronchi et al., 2021). Despite the unsurpassed advantage of HD-MEAs recordings, the high volume and intercoupling of signals on neighboring channels remains a considerable challenge in the analysis and accurate online detection of single-unit activity. A commonly used method to quantify neural activity in MEAs is adaptive threshold crossing spike detection. In this method, the spike detection threshold is set as a multiple of the signal noise, which is commonly estimated with the Root-Mean-Square (RMS) of the signal (Ronchi et al., 2021; Sommer et al., 2022; Miyahara et al., 2023; Bertacchi et al., 2024; Vinci et al., 2025; Setsu et al., 2025; Kesdoğan, et al., 2024; Fenton et al., 2024; Hoang et al., 2025; Zhao et al., 2025). Additionally, to account for changes in noise conditions the threshold can be repeatedly estimated from short windows of the signal. Unfortunately, the adaptive RMS estimate lacks sensitivity during periods of high network bursting activity, which are frequently observed in mature neurons *in vitro* (Murphy et al., 1992; Wong et al., 1993; Rhoades & Gross, 1994; Kamioka et al., 1996; Canepari et al., 1997; Voigt et al., 1997; Segev et al., 2001; Harris et al., 2002; Wagenaar et al., 2006; Chiappalone et al., 2007; Chen and Dzakpasu, 2010; Colombi et al., 2013; Suresh et al., 2016). Other estimators for the signal noise, such as the Median of Absolute Deviations (MED), can be used in threshold crossing techniques to address the frequency dependence of the conventional RMS method (Quiroga et al., 2004). Beyond simple spike detection, commonly used methods include spike sorters, such as Kilosort4 (Pachitariu et al. 2024). These methods typically involve template-matching and clustering steps to separate detected spikes into identified units. However, these steps incur time-consuming computations that are exacerbated by dense arrays and high activity. Thus, faster and accurate spike detection methods that can accommodate fluctuating activity are desirable for HD-MEA data.

Given the many advantages of multi-electrode arrays, they have been commonly used for drug screening (Stett et al., 2003; Natarajan et al, 2011; Valdivia et al., 2014; Hondebrink et al., 2016; Passaro et al., 2021; Ronchi et al., 2021;Sommer et al., 2022;Lee et al., 2022; Kim et al., 2023; Ganbat et al., 2023; Kesdoğan, et al., 2024;Lee et al., 2024; Zhao et al., 2025; Fofie et al., 2025). Drug screening is essential for the development of new therapeutic approaches to treat a wide range of neurological disorders (Cui et al., 2006; Kaltenbach et al., 2010; Hill et al., 2010; Kitaguchi et al., 2017; Kouroupi et al., 2020; Silva and Haggarty, 2020; Ganbat et al., 2023; Fofie et al., 2025). Considering the costs involved in drug development and the significant public health implications, establishing high quality pipelines for the fast and accurate interpretation of HD-MEA results is critical.

Here we describe a pipeline for accurate and fast detection of spikes in HD-MEA recordings and tested it in conditions of high neuronal activity during bursting behavior, and enhanced activity elicited by the application of a pharmacological agent. Using the MED method, we consistently detected ∼50% of spikes that were missed by the RMS method during bursts. Furthermore, the subcellular spatial resolution of the HD-MEA leads to the detection of action potentials by more than one electrode, resulting in many duplicate detections. Therefore, we developed a heuristic approach to quickly de-duplicate spike detections found by the MED method (DP-MED). We compared our DP-MED algorithm to a commonly used sorting algorithm, KS4, using simulated data from model neurons. Overall, DP-MED shows more precision than KS4 across a range of increasing firing rates, but it shows less accuracy than KS4 in this range. Thus, both methods exhibit comparable performance in spike detection. Last, we found no differences between these methods in assessing the number of spikes when neuronal spiking was enhanced in the presence of the cholinergic agonist carbachol. However, the analysis in DP-MED was 40-fold less time-consuming compared to KS4. These results suggest that the pipeline we developed is less computationally expensive and comparable in performance to commonly used sorting algorithms like KS4. The DP-MED method is a suitable and precise tool for drug screening in neuronal cultures that commonly show bursting periods during development and maturity.

## 2. Methods

### 2.1. Primary cortical culture preparation and data recording

#### 2.1.1. Cell Culture

Primary neuronal cultures were prepared from commercially available rat cortical neurons (ThermoFisher, A1084001), using standard protocols (Heiney et al., 2022; Genocchi et al., 2024). Primary rat cortical astrocytes (Cell Applications, R882A-05n), were plated, expanded and stored in Liquid Nitrogen. On the day of plating, astrocytes were thawed and combined with the neurons (ratio 1:8; astrocyte/neurons), following the provider’s protocol. Neurons were thawed and resuspended to a concentration of ∼3,000,000 cells/mL. Droplets of ∼80 uL containing 250,000 neurons were plated either on HD-MEAs or 12 mm coverslips, to conduct immunohistochemistry, morphological analysis and whole-cell recordings. Before plating, the HD-MEAs were treated with TergAzyme to increase the hydrophilicity of the surface, followed by primary coating with poly-D-lysine (0.1 mg/ml; 1-3 hours inside of the incubator) and secondary coating with b-laminin (0.02 mg/ml; 1 hour inside of the incubator) following MaxWell Biosystems coating protocol.

#### 2.1.2. Immunohistochemistry

Coverslips, or HD-MEAs, were fixed with 4% paraformaldehyde (warmed at 37C) for 15 min, followed by three 10-min washes in PBS. Blocking was performed using 5% donkey serum in 0.3% Triton-PBS-1X for 1 hr. Primary antibodies, including MAP2 (anti-chicken, 1:500) for neuronal markers and GFAP (anti-rabbit, 1:500) for astrocytic markers, were incubated for 2 hrs. at room temperature. After three additional 10-min PBS washes, the cells were incubated for 2 hrs. in the dark with the corresponding secondary antibodies; Alexa 488 (donkey anti-mouse, 1:500) and Alexa 594 (donkey anti-rabbit, 1:500). Finally, after three 10-min PBS-1X washes, the coverslips were mounted with Fluoromount-DAPI. Confocal images were taken in Leica Stellaris 8 FALCON using the 40X objective. Purchase of the Leica Stellaris 8 FALCON was supported by Award Number 1S10ODO34260 from the National Institute of Health.

#### 2.1.3. Electrophysiology Patch Clamp

Neurons were recorded at room temperature in current clamp mode using a SutterPatch digital amplifier. Cells were bathed in artificial cerebrospinal fluid (ACSF) solution with the following composition (in mM): 125 NaCl, 25 NaHCO3, 1.25 NaH2PO4, 3 KCl, 2 CaCl2, 1 MgCl2, 3 myo-inositol, 0.3 ascorbic acid, 2 Na-pyruvate, and 15 glucose; continuously oxygenated with 95% O2 and 5% CO2. Coverslips containing the neurons were transferred to the recording chamber mounted on an Olympus BX51 W1 microscope. Neurons were visualized using 40X objectives and DIC illumination. The internal solution in the recording pipette had the following composition (in mM): 120 K-gluconate, 10 Na-gluconate, 4 NaCl, 10 HEPES-K, 10 Na phosphocreatine, 2 Na-ATP, 4 Mg-ATP, and 0.3 GTP, adjusted to pH 7.3 with KOH. In a subset of experiments, Alexa-488 (20 μM, Invitrogen) was included in the internal solution for *post hoc* morphologic reconstruction of the recorded neurons. The recording pipettes were pulled from thick-wall borosilicate glass capillaries using a horizontal puller (P-97, Sutter Instrument) and had a resistance of 3-8 MOhm.

#### 2.1.4. Electrophysiology HD-MEA

Extracellular signals were recorded using the MaxOne recording unit with single-well HD-MEAs (MaxWell Biosystems). The HD-MEAs have 1,024 recording channels which can be routed to 1,020 of 26,400 platinum electrodes arranged in a 17.5 um pitch grid of 120x220 electrodes. The electrodes are coated in platinum-black by the vendor (MaxWell Biosystems) to reduce impedance and improve recording characteristics. Each electrode is 5 μm x7 μm.

Recordings were performed every other day (starting on day *in vitro* (DIV) 5). To identify electrodes that can record neuronal activity, we configured the array to record from columns that are sequentially moved across the array. After scanning the activity of the whole array, ∼1,000 electrodes of interest were selected for recording based on their observed firing rate and spike amplitudes. These were subsequently simultaneously recorded for 15 minutes. This approach allowed us to observe network activity in our cultures.

#### 2.1.5. Drugs and Pharmacology

All chemicals were obtained from Sigma Millipore. Carbachol (30 µM) was applied to the bath using a micropipette after 1 min of baseline recording. The neuronal activity was then recorded for an additional 15 min. In the control condition, an equal volume of sucrose solution (30 µM, matched for osmolarity) was used.

### 2.2. Spike Detection Methods and Data processing

#### 2.2.1. Spike Detection

Raw signals from the MEAs were high pass filtered at 300 Hz with a 3rd order Butterworth filter. A forward and backward pass over the data was performed to achieve zero-phase filtering. After this, the noise in each channel was estimated in 0.1 s segments using two methods. The first is the Root-Mean-Square method (RMS), where the RMS of the mean-subtracted voltage trace is used to estimate the noise standard deviation. The second method is the median method, where a scaled median of absolute deviations is used to estimate the noise as follows:

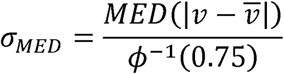

where *φ*^-1^ is the inverse standard cumulative density function (CDF) and *v* is a voltage trace from one HD-MEA channel. Under the assumption of Gaussian noise, the normal CDF factor scales the median to be an estimate of the noise standard deviation, where *φ*^-1^(0.75) is the median of the distribution |*X*|, *X* ∼ Normal (0,1)). This estimation method is less sensitive to outliers in the voltage signal, the main source of which is spikes (Quiroga et al., 2004). After noise estimates were calculated for each channel, spike detection thresholds were calculated as 7 times the noise estimate. Extracellular action potentials (EAP) have large negative excursions, so spikes times were detected as the minimum voltage following a negative threshold crossing. Additional threshold crossings within 1ms were discarded to avoid multiple detections of the same spike.

#### 2.2.2. Burstiness

Following Wagenaar et al. 2005, we define the burstiness of a recording to be the proportion of spikes contained in the top 15% of time bins.

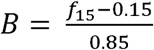

Where f_15_ is the fraction of spikes contained in the top 15% most active time bins. If spikes are uniformly distributed throughout the recording, then the top 15% of bins will contain 15% of the spikes, and the metric will be 0. If the top 15% contains all spikes, the metric will be 1. We report this metric using time bins of 0.5s.

#### 2.2.3. Burst Detection

Synchronized volleys of spikes across many cells occur at regular intervals in dissociated cultures (Wagenaar et al., 2006). To detect these events, we used the ISI-N detection algorithm. This method computes the inter-spike-interval between N spikes (called ISI-N) and detects a burst when the ISI-N drops below a set threshold (Bakkum et al. 2014). This method has the advantage that no smoothing of network firing rates is needed, so ISI-N threshold crossings coincide with the first and last spike times of the detected burst. In this paper, we used an N-value based on observing activity in half of the channels, filtering out the channels with less than 0.1 Hz spike activity. The values of N range from ∼100 to ∼500 across different cultures.

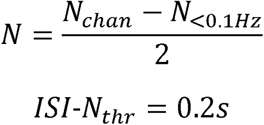

where *N_chan_* is the number of channels and *N*_<0.1*Hz*_ is the number of channels with less than 0.1 Hz observed spike rate.

#### 2.2.4. Spike De-duplication

For each detected extracellular action potential (EAP), we look in a spatiotemporal neighborhood for other possible detections of the same spike. Here we use a circle of radius 50 um and a time window of 2 ms for consideration of duplicate detections. These dimensions match the typical extracellular action potential spatial and temporal extent that we observe in our system (Supp. Fig. 4). For all spikes within this neighborhood, we only retain the one with the largest amplitude. This can be thought of as the “center” of the EAP. This method is a fast and approximate way to filter out multiple observations of the same action potential without attempting to sort spikes.

#### 2.2.5. Kilosort 4

KS4 is an open-source spike sorting method designed for *in vivo*, multi-electrode silicon probe recordings, but it can also be used for high-density *in vitro* arrays (Pachitariu et al., 2024). Briefly, KS4 uses a template matching approach to detect spikes. Each template is a multi-channel waveform that represents the typical waveform of a given neuronal unit. They are found via clustering of detected multi-channel waveforms. The initial estimation of neuronal units is used in a second pass over the data for more sensitive spike detection and improved clustering. The output of KS4 is therefore a set of neuronal units, each with their own multi-channel spike template, and spike times for each of the units. This is different to the RMS and MED methods, which associate a spike with a channel and not with a neuronal unit. Our choice of parameters for KS4 was based on the HD-MEA properties and bursting behavior of the cultures.

#### 2.2.6. Data Simulation

Simulated High-Density Microelectrode Array data was generated using the LFPy and MEArec python packages (Lindén et al., 2014; Buccino & Einevoll, 2021). A 10 x 10 electrode array with 15 um electrode pitch was created, and cells were placed on it at a density of ∼2500 cells / mm^2^. Extracellular waveforms were generated from LFPy by first calculating transmembrane currents from multi-compartmental models of reconstructed cortical cells. The transmembrane currents are used in the Line Source Approximation to determine the EAP from any observation point (Lindén et al., 2014), thus generating an EAP for any electrode array. After cells were placed and EAPs were generated, network activity was simulated with Poisson firing statistics for each cell. Network burst occurrences were modelled as a Poisson process, and their duration was drawn from a uniform distribution between 200 and 300 ms. All neurons had elevated firing rates for the duration of the burst, after which they returned to their basal firing rate.

#### 2.2.7. Spike detection method comparison

For RMS and MED detectors, we can simply check the highest amplitude channel of a waveform for detection at each spike time. However, for spike sorted data it is not guaranteed that the detected unit waveforms correspond to ground truth waveforms because spike sorters may falsely merge or split units, impacting the estimate of a unit’s waveform. This complication motivated three evaluation methods for spike sorted data, iii) Spike matching. For each spike in the ground truth data, look for a matching detection in the spike sorted data, irrespective of the unit label. This is the most liberal of the three methods. (ii) For each unit in the spike sorted data *j_S_*, find the best matching ground truth spike train *i_G_* and assign *j_S_* to *i_G_*. The spikes in *j_S_*are then evaluated against those in *i_G_*. (iii) Template matching. Match the spike sorted units to the ground truth units based on the cosine similarity of the identified spike waveforms to the ground truth. Data is not shown for methods (i) and (ii); we chose (iii) template matching as is the most conservative option. Figure 3B-D shows the results for the template-matched assessment.

### 2.3. Statistical Analysis and benchmark evaluation

#### 2.3.1. Statistical Analysis

All statistical analyses were conducted using Python v.3.10.2. All levels of statistical significance were set at \**P* < 0.05, **P<0.01 and ***P<0.001. Shapiro–Wilk normality test was conducted to evaluate the distribution of the data and select the corresponding statistical analysis. All statistical analyses are reported in the results and figure legends. To test the development of cultures, we performed a terminal segment test of the activity and firing rate curves using an asymptotic slope test (Fig. 1*C*). For each dataset, the curve was interpolated into a uniform x axis, and the final 20% of the points were extracted. A simple linear regression was then fitted to this terminal segment. Plateau onset was operationally defined as a terminal slope smaller than 0.05 units (Slope<0.05), indicating negligible change over time.

**Figure 1.**
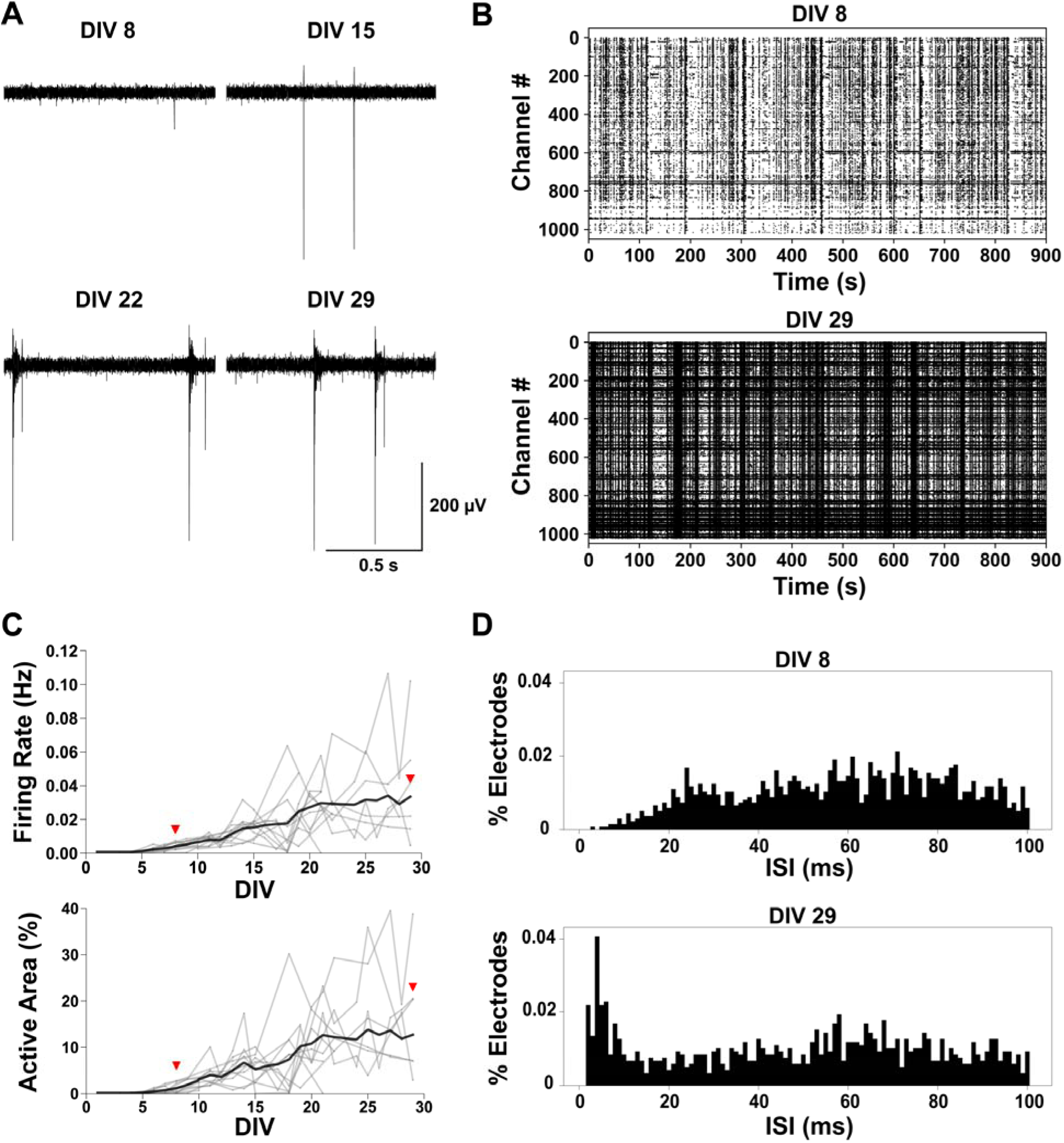
Characterization of primary cortical cultures development in HD-MEAs. **A.** Filtered voltage traces (300 Hz high-pass) of a single channel showing the evolution of spikes detected in 1 second at 4 different DIVs 8, 15, 22 and 29. Scale Bar is 200 μV and 0.5 s. **B.** Raster plots from a representative culture showing independent spikes (dots) detected across 15 minutes of network recording for DIV 8 and 29 (∼1000 channels at the same time). **C.** Top, number of electrodes that detected activity represented as active area percentage across DIV for primary cortical cultures. We observed the first signs of sparse activity around DIV 4 and there was an overall increase in activity until ∼DIV 20, reaching a plateau after three weeks *in vitro.* Bottom, average firing rate for the whole HD-MEA across DIV for primary cortical cultures. There was an overall increase in activity until ∼DIV 20, reaching a plateau after three weeks *in vitro*. Every independent culture is represented as a gray line, and the averages are shown as a black line in each plot. Red triangles point out the average culture activity at DIV 8 and 29 (n=11 cultures). **D.** Histograms of percentage of electrodes showing activity in a particular inter-spike interval (ms) at DIV 8 and 29. We observed slow activity at DIV8 and a change in activity distribution at DIV29 with more channels detecting events more frequently (n=11 cultures).

#### 2.3.2. Detector Runtime Measurements

Runtime of the spike detection algorithms used in this paper were taken on a lab workstation featuring an Intel Xeon w5-3423 12 Core CPU, NVIDIA RTX 5000 GPU, 128GB DDR5 RAM, and a 4TB HDD. Runtimes are reported as the real time needed to process HD-MEA data on this system. Recordings were all within a few seconds of 15 minutes in duration and contained approximately 1000 electrode channels.

##### Precision, Recall, and Accuracy

For the simulated data we used standard definitions for; precision, recall, and accuracy, which are usually used to quantify detector performance:

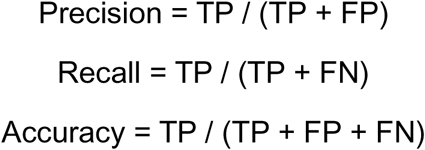

Where TP is True Positive, FP is False Positive, and FN is False Negative.

## 3. Results

### 3.1. Development and maturation of cortical networks

We established long-term cortical neuron cultures to study network dynamics using HD-MEAs. To this extent, we record neuronal activity across the entire active area of the HD-MEA at different days *in vitro* (DIV) (see Methods). Figure 1*A* shows the evolution of the same electrode’s activity across DIVs 8, 15, 22, and 29. There is an increase in firing rate between DIV8 and DIV22, reaching a plateau after three weeks (Fig. 1*A*; DIV8, 0.09 ± 0.04 Hz; DIV15, 0.7 ± 0.2 Hz; DIV22, 4.06 ± 0.8; Hz; DIV29, 3.96 ± 0.6 Hz; n=14 individual channels). After examining the average activity of the HD-MEA, we selected approximately 1000 electrodes to characterize the network activity (see methods). The selected individual channels showed higher amplitude spikes and higher firing rates, using thresholds of ∼50 μV for spike amplitude and ∼1.0 Hz for firing rates. As shown in Figure 1*B*, there is an increase in overall activity across the entire chip between DIV8 and DIV29, similar to what we observed in individual channels (Fig. 1*B*; DIV8, 0.004 ± 0.001 Hz; DIV29, 0.034 ± 0.008, n=11). A complete timeline analysis of the cultures maturation in the HD-MEAs showed that the number of electrodes detecting activity across the entire active area increased from DIV5 to DIV22, reaching a plateau after three weeks *in vitro* (Fig. 1*C*, top; DIV8, 0.94±0.31%; DIV15, 4.91±0.92%; DIV22, 12.19±2.17%; DIV29, 12.71±3.11%; Asymptotic Slope Test, plateau slope: 0.00913; n=11). This is consistent with the average firing rate of neurons in the network, which follows a similar trajectory (Fig. 1*C*, bottom; DIV8, 0.004 ± 0.001 Hz; DIV15, 0.015 ± 0.002 Hz; DIV22, 0.029 ± 0.005 Hz; DIV29, 0.034 ± 0.008 Hz; Asymptotic Slope Test, plateau slope: 0.00047; n=11). The analysis of all active channels on the chip showed a similar pattern, with an increase in the percentage of channels exhibiting shorter inter-spike intervals (ISIs) after three weeks *in vitro* (Fig. 1*D*; DIV8, ISI distribution, 5.16±0.13 ms; DIV29, ISI distribution 4.06±0.12 ms; Two-sample Kolmogorov-Smirnov test; *p<0.01*). In parallel experiments, we assessed the cellular composition of cultures grown on coverslips at similar DIVs using immunostaining (Supp. Fig. 1*A*). Three weeks post-plating, immunostaining for the microtubule-associated protein 2 (MAP2) indicated that approximately 25% of the cells were neurons, while immunostaining for the intermediate filament-III protein (GFAP) revealed approximately 75% astrocytes. In addition, immunostaining with VGLUT1, a vesicular glutamate transporter, and a GAD67 antibody, which labels an enzyme isoform responsible for GABA synthesis, showed abundant excitatory and inhibitory neurons, respectively (Supp. Figs. 1*B* and *C*). These results indicate that at 3 weeks network activity arises from an interconnected network consisting of mature excitatory and inhibitory neurons. Accordingly, current-clamp recordings three weeks post-plating, showed characteristic responses of mature neurons and abundant synaptic activity (arrows), reflecting extensive connectivity (Supp. Fig. 1*D–E*). Sholl analysis indicated extensive dendritic branching near the soma and fewer projections farther away. An average of 12 intersections were observed within 20-60 μm from the soma, which decreased with distance (Supp. Fig. 1*F*). Overall, these results indicate that our cultures reach maturity and stability in basic firing properties after three weeks *in vitro*. Furthermore, our co-cultures of neurons and astrocytes contain both excitatory and inhibitory neurons, exhibiting normal network connectivity which is consistent with prior studies (Björklund et al., 2010; Beaudoin et al., 2012, Ito et al., 2013; Harrill et al., 2015).

As shown in Figure 1*B*, network activity increased after three weeks *in vitro*, with spontaneous firing and prominent bands of synchronized activity. These bands of synchronized activity are roughly periodic (Supp. Fig. 2*A*), with periods ranging from a few seconds to a few minutes depending on the age of the culture (Supp. Fig. 2*B*) and is consistent with network bursts previously described in cortical cultures (Kamioka et al., 1996; Canepari et al., 1997; Voigt et al., 1997; Wagenaar et al., 2005; Chiappalone et al., 2007). Based on features of the ISI-N distribution analysis (see Methods) performed on our data (data not shown), we define a network burst as a period of activity in which all inter-spike intervals between the *i*^th^ spike and the (*i* + *N*)^th^ spike are 200 ms or less (Supp. Fig. 2*A*). In agreement with previous studies (Wagenaar et al., 2005; Chiappalone et al., 2007), network bursting frequency increased as the cultures matured, reaching a plateau after three weeks, as reflected by an increase in burst frequency (Supp. Fig. 2*B*, top; DIV6, 0.005 ± 0.0008 Hz, n = 7; DIV15, 0.11 ± 0.03 Hz, n = 10; DIV22, 0.21 ± 0.04 Hz, n = 7; DIV29, 0.17 ± 0.04 Hz, n = 6). In addition, we observed an increase in the number of spikes per burst (Supp. Fig. 2*B*, bottom; DIV6, 2,168 ± 479 spikes, n = 7; DIV15, 4,173 ± 856 spikes, n = 10; DIV22, 8,537 ± 1,014 spikes, n = 7; DIV29, 6127 ± 1063 spikes n = 6). Overall, these data indicate that the cultures reach a stable level of activity around three weeks *in vitro*; therefore, most of our studies were conducted at this time point.

### 3.2. Performance of Spike Detection Methods

We next compared the performance of the RMS and MED methods in the analysis of spontaneous network activity, at three weeks *in vitro* (Fig. 2). As shown above, at this stage, network activity exhibits periods of low spontaneous activity interrupted by periods of highly synchronized activity or bursting (Fig. 2*A*). At low frequencies, both the RMS and MED algorithms exhibit similar performance (Fig. 2*A*, bottom left), but the MED method detects 19.70% ± 0.319% more spikes than the RMS method (Fig. 2*A-B*, left; n=50 recordings from 6 cultures). However, during periods of high network bursting, there was a dramatic difference between the two methods (Fig. 2*A-B*, right); MED detected nearly twice as many spikes as the RMS method; of all detected spikes, 49.56% ± 0.26% were only detected by MED (Fig. 2*B*, right; n=49 recordings from 6 cultures). We also characterized the differences between the algorithms as a function of the burstiness of a recording session (Supp. Fig. 3). The discrepancy between the detectors grows as the burstiness of a recording session increases (Supp. Fig. 3*B-C*). Indeed, the percentage of spikes only found in MED is positively correlated with the burstiness of the recording (Supp. Fig. *3B-C*; One-way ANOVA, *p<0.001)*. The lower performance of the RMS method in spike detection under both low and high neuronal activity prompted us to optimize the MED method to achieve reliable spike detection in our cultures, particularly during network bursts.

**Figure 2.**
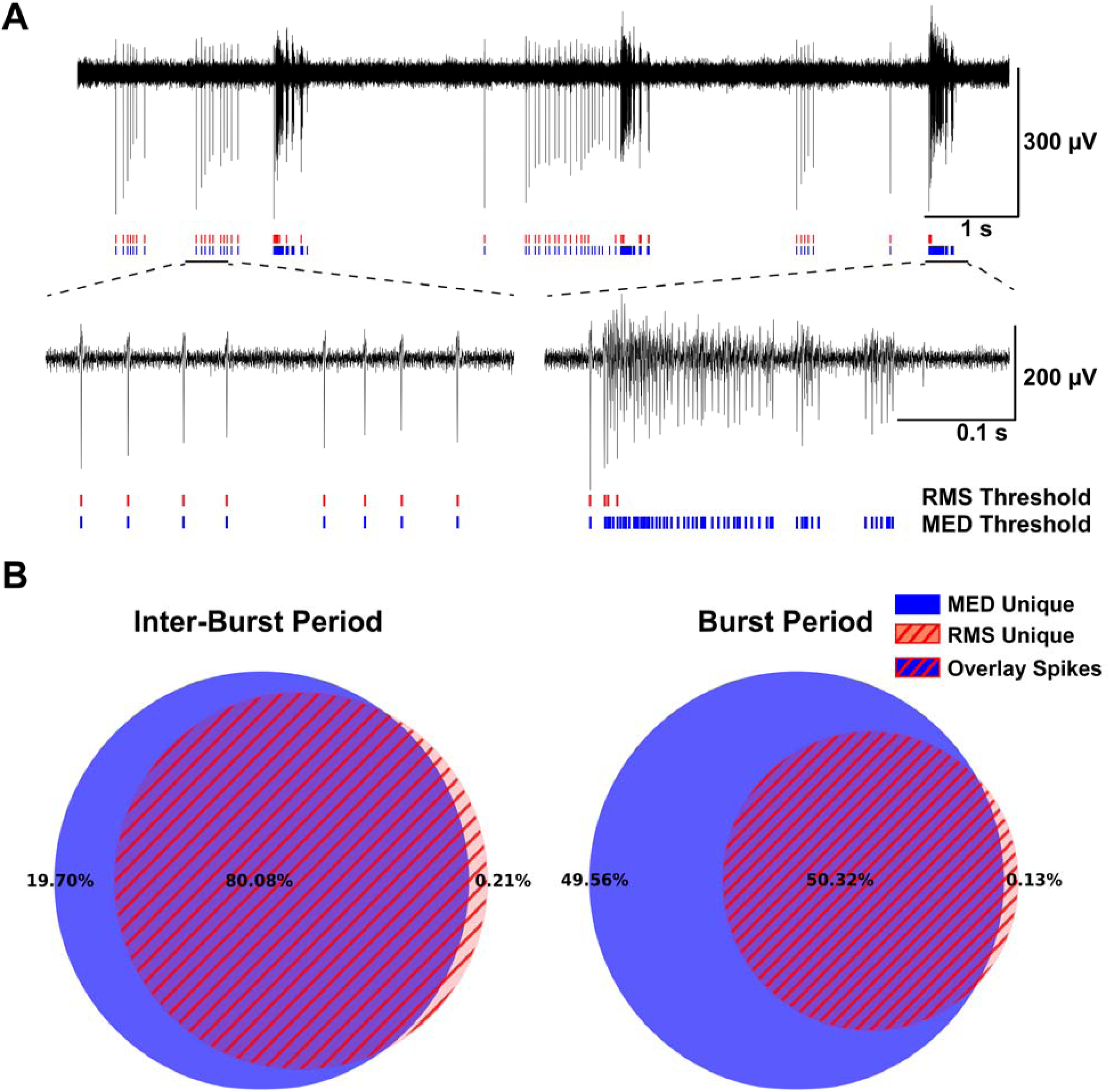
Spike detection method comparison. **A.** Spike detection example, 10 s extracellular voltage trace showing low frequency spontaneous activity and high frequency network bursts. Top Scale Bar is 300 μV and 1 s **B.** Overlap of detected spikes in all recordings. Left: The percentage of spikes unique to each detection method is plotted in a Venn diagram for inter-burst periods (not to scale). Higher percentages in the middle of the diagram indicate more overlap between detection methods. Right: The percentage of spikes unique to each detection method is plotted in a Venn diagram for burst periods (not to scale). Lower percentages in the middle of the diagram indicate less overlap between detection methods (N=50 recordings, 6 cultures). Scale Bar is 200 μV and 0.1 s.

### 3.3. Performance of DP-MED and KiloSort-4 with simulated data

At three weeks *in vitro* neurons exhibited extensive processes and connectivity (Supp. Fig. 1), therefore, we expected that the activity from a single neuron may be detected by multiple electrodes. We estimated that approximately 38% of spikes detected by each method are duplicates, appearing as lower-amplitude action potentials in neighboring electrodes (MED 38.78%±2.1% of detections duplicated, n=50; RMS 37.98%±2.1% of detections duplicated, n=50 recordings from 6 cultures). To resolve this issue, we implemented a spike de-duplication (DP) step for the MED method, which considers the timing, spatial distribution, and amplitude of each event (Supp. Fig. 4). To test the efficacy of our DP algorithm, we compared the combined DP-MED method to KS4 (Pachitariu et al., 2024). To accomplish this, we simulated data that resembles our cortical recordings, including periods of low spontaneous firing and highly synchronized network bursts. We recreated a small network of 25 pyramidal neurons on a 100-channel array (Fig. 3*A*). The simulated neurons had similar firing patterns, with background spontaneous activity of 0.1 Hz, which was elevated to increasing frequencies, in the 1Hz to 300 Hz range, during bursting periods (Fig. 3*B-D*). Further analysis indicated that the precision of spike detection is highly sensitive to firing rate in both methods (Fig. 3*B*; Two-way ANOVA, Frequency, *p<0.001*, detection method effect, *p<0.001;* interaction, *p <0.001*). Interestingly, DP-MED is significantly more precise than KS4 at high firing rates (Fig. 3*B*; Tukey post-hoc, e.g. 1 Hz, *p>0.05,* 25 Hz *p<0.05* and 300 Hz *p<0.001;* n=5). This indicates that KS4 detects more false positives than DP-MED. Similar to the precision results, recall decreases as firing rate increases in both detection methods (Fig. 3*C*; Two-way ANOVA, Frequency, *p<0.001*, detection method effect, *p<0.001;* interaction, *p <0.001*). However, KS4 shows better recall than DP-MED at different firing rates (Fig. 3*C*; Tukey post-hoc, e.g. 1 Hz and 25 Hz, *p>0.05;* 10 Hz, *p<0.05 and* 300 Hz *p<0.001;* n=5). This means that KS4 detects fewer false negatives than DP-MED. Finally, consistent with the previous results, accuracy decreases with increasing firing rates in both detection methods (Fig. 3*D*; Two-way ANOVA, Frequency, *p<0.001*, detection method, *p<0.01*, interaction, *p<0.05*). KS4 is significantly more accurate than DP-MED but only at high firing rates (Fig 3*D*; Tukey post-hoc, e.g. 1 Hz and 25 Hz, *p>0.05;* 200 Hz, *p<0.05* and 300 Hz, *p<0.01;* n=5). This indicates that at high firing rates KS4 detects more true positives, fewer false negatives, or both, compared with DP-MED. Overall, this demonstrates a tradeoff between recall and precision for KS4 and DP-MED, with KS4 finding more spikes but more false positives and DP-MED finding fewer spikes but fewer false positives.

**Figure 3.**
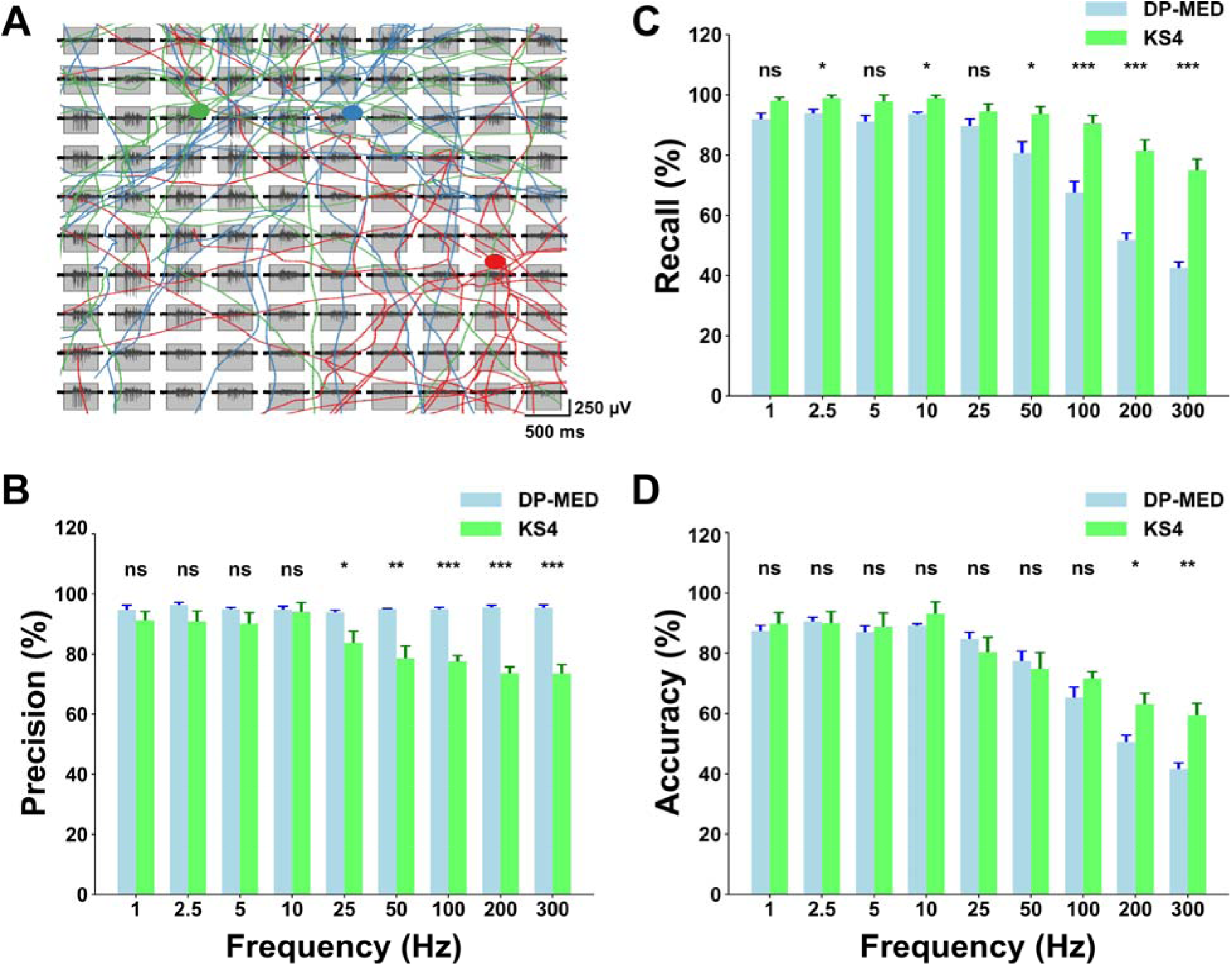
Performance of different detection methods in simulated data. **A.** Example traces from a simulated burst on a 10 x 10 electrode HD-MEA. Superimposed on the traces we show a morphological representation of 3 simulated cells (green, red and blue). Scale Bar is 250 μV and 500 ms. (**B-D**) Precision, recall and accuracy comparisons for DP-MED and KS4 detection methods across within-burst firing rates. Note the steep drop-off in accuracy and recall for DP-MED when firing frequencies become high. **B.** DP-MED is significantly more precise than KS4 at high firing rates (Tukey post-hoc, 1-10 Hz, *p>0.05* and 25 Hz-300 Hz, *p=0.0306, 0.0043, 0.0001, 0.0001, 0.0001;* simulated traces). **C.** KS4 shows better recall than DP-MED across different firing rates (Tukey post-hoc, 1 Hz, 5 Hz and 25 Hz, *p>0.05*, 2,5Hz, *p= 0.02*, 10 Hz, *p= 0.003*, 50 Hz-300 Hz, *p=0.02, 0.0007, 0.0001, 0.0001;* n=5 simulated traces). **D.** KS4 is significantly more accurate than DP-MED at high firing rates (Tukey post-hoc, 1-100 Hz, *p>0.05,* 200 Hz-300 Hz, *p=0.018, 0.0045;* simulated traces).

### 3.4. Performance of DP-MED and KiloSort-4 with high volume data from cortical cultures

We next compared the performance of both detection methods under high neuronal activity induced by a pharmacological agent, as is commonly done in drug screening trials. Previous studies have shown functional cholinergic transmission in cortical primary cultures (Wang et al. 1994; Tateno et al. 2005a; Tateno et al. 2005b; Hammond et al. 2013). We therefore stimulated the cultures with carbachol, a non-selective cholinergic receptor agonist. As shown in figure 4*A*, the addition of carbachol increased spontaneous activity and bursting patterns at individual channels and across the entire chip (Fig. 4*A-B*, control(blue) vs carbachol (red); cumulative distribution probability of individual electrodes firing rate, 0.53±0.4; 1.75±1.5; Two-sample Kolmogorov-Smirnov test; *p<0.001*; inter-burst interval, 7.68±0.32; 2.18±0.57 (s); Mann-Whitney U; *p< 0.05;* n=4). We measured the number of spikes detected by both DP-MED and KS4 under control conditions and after carbachol application. Consistent with our findings in simulated data, we observed no significant difference in the number of spikes detected by both methods under control (Fig. 4*C*; 559.86±12.83; 1130.849±22.31; Mann-Whitney U; *p>0.05;* n=4) or after carbachol stimulation (Fig. 4*D*; 1785.51±49.81; 2702.25±53.96; Mann-Whitney U; *p>0.05;* n=4). Nevertheless, KS4 tended to detect more spikes than DP-MED (Fig. 4*C-D*), which is consistent with the observations of true and false positives from our simulated data. These results indicate that DP-MED is comparable to KS4 regarding the number of spikes detected in a drug screening experiment. However, despite both methods’ similar performance in spike detection, KS4 is time-consuming and its processing time increases dramatically as spike density in the recordings rises. To compare the computational speed of DP-MED, we calculated the average time each method required to process the same batch of HD-MEA recordings (∼1,000 channels over 900 seconds at 20 kHz). The processing time is significantly lower in DP-MED compared to KS4, approximately 40 times faster (Fig. 4*E*; 0.065±0.003; 2.69±0.129; Mann-Whitney U; *p<0.001;* n=54). Overall, the two methods perform better in different contexts, but their detection performance is comparable. DP-MED is more precise but misses some spikes that KS4 detects, whereas KS4 detects more spikes but produces more false positives than DP-MED. Thus, a major advantage of DP-MED over KS4 is that it is less time-consuming.

**Figure 4.**
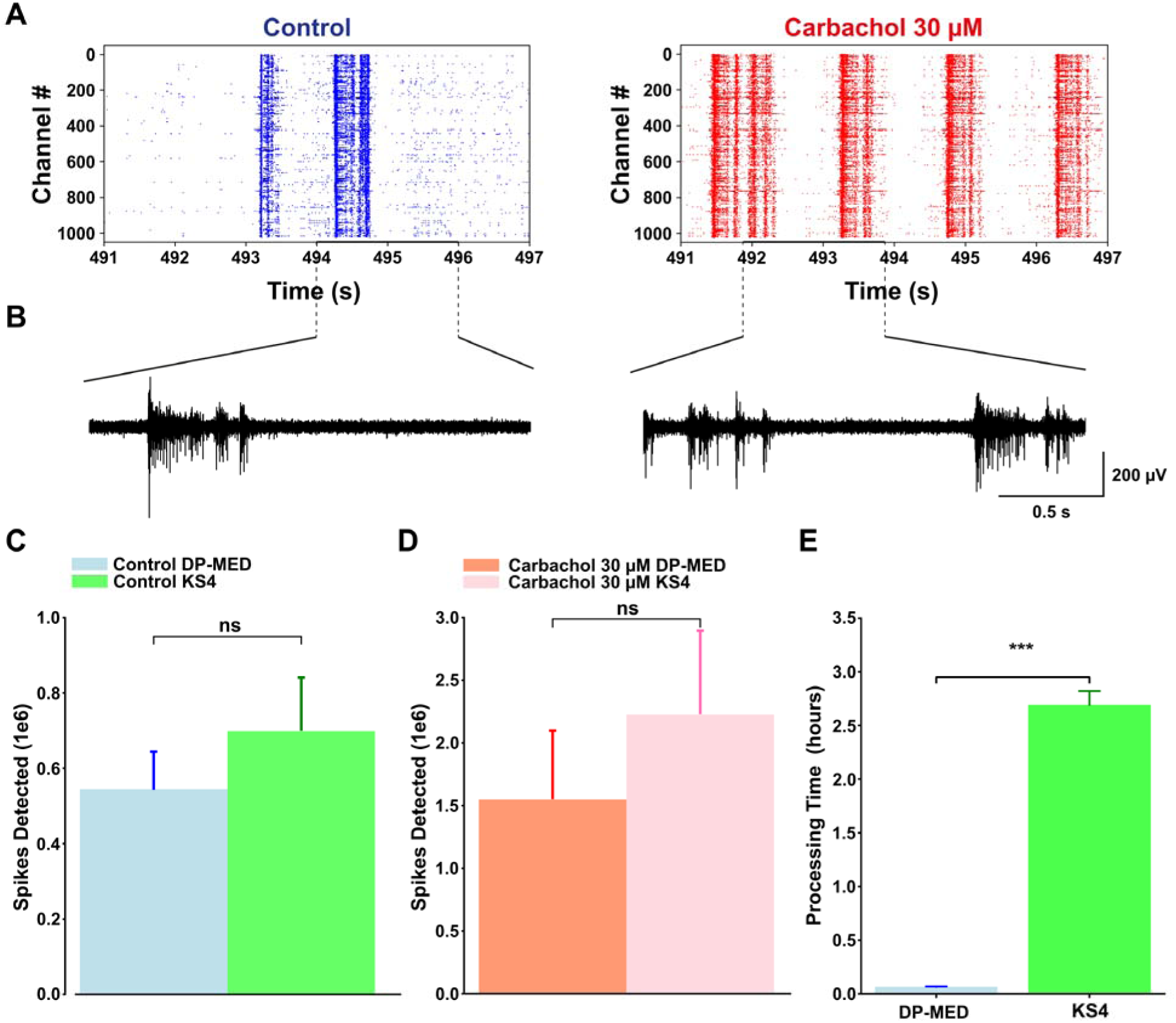
Spike detection method comparison in a pharmacology experiment. **A**. left: representative raster plot showing spikes detected (dots) across all channels for control condition (blue). Spikes were identified using the MED threshold detection method. Right: raster plot showing spikes detected (dots) across all channels after the application of 30 µM carbachol (red), also using the MED threshold detection method. **B.** Filtered voltage traces (300 Hz high-pass) showing network bursts as recorded by the electrodes over a 2-second window (highlighted in black). Scale Bar is 200 μV and 0.5 s. **C.** Number of spikes detected by both methods in control condition. The Mann-Whitney U test revealed that de-duplicated MED (light blue) and KS4 (green) detected a similar number of spikes. **D**. Average of spikes detected by both methods following carbachol application in every culture. No significant difference was observed between the number of spikes detected by the de-duplicated MED (red) and KS4 (pink). Across both methods, spike counts were higher after carbachol application compared to control conditions (n=4 independent cultures). **E.** Runtime of the spike detection algorithms used in this paper. n=55 recordings of 15 minutes in duration and 1000 channels (6 cultures). All algorithms were run on a machine with an Intel Xeon w5-3423 12 Core CPU, NVIDIA RTX 5000, 128GB DDR5 RAM, and a 4TB HDD.

## 4. Discussion

We designed a fast, simple spike detection pipeline that combines the MED method with a heuristic spike de-duplication step. Compared to RMS-based threshold detectors, our method, which we termed DP-MED, provides more accurate spike counts during high frequency firing, as observed in spontaneous network bursts and high network activity induced by a pharmacological agent. Across several analytical parameters used to characterize network activity the DP-MED method is comparable in performance to the KS4 sorting algorithm. However, in analyzing large HD-MEA datasets, DP-MED was on average, 40 times faster than KS4 when run on a CPU.

Based on the developmental profile of individual active electrodes, overall firing rate and network bursts, our cortical cultures reach maturity after three weeks *in vitro;* this is consistent with the studies that have used cortical cultures in MEA recordings (Chiappalone et al., 2004; Chiappalone et al., 2005; Wagenaar et al, 2006). At this stage of development, the synaptic mechanisms that control network dynamics such as synapse formation and dendritic branching (Massengill et al., 1997; Fischer et al., 2000; Lesuisse & Martin, 2002; Kato-Negishi et al., 2004; Ito et al., 2013; Harrill et al., 2015), as well as biochemical markers of glutamate metabolism have stabilized (Björklund et al., 2010). In parallel coverslip experiments, we showed that our cultures contained excitatory and inhibitory neurons, a cellular composition expected for cortical primary cultures (Björklund et al., 2010; Beaudoin et al., 2012, Ito et al., 2013; Harrill et al., 2015). Consistent with the culture maturation measurements in HD-MEAs, our cultures also showed normal electrophysiological properties, including excitatory postsynaptic potentials. This suggests that after three weeks in vitro, the cultures are sufficiently mature to allow characterization of network properties.

Traditionally, bursts are defined as periods of highly synchronized activity in unperturbed networks widely observed in dissociated cortical cultures (Wagenaar et al., 2006). However, in some of our recordings, pharmacological manipulation causes the network activity to oscillate slowly rather than burst intensely, as observed in control trials (data not shown). In these situations, the definition of burst may be complicated, making the interpretation of data more difficult. Furthermore, ISI-N–based burst detection is directly influenced by the number of spikes detected, and the parameters must be normalized to account for this (see *Methods*). Our results show that the performance of the RMS-based method declines considerably more during network burst periods than during inter-burst intervals. In contrast, MED maintains stable performance under these conditions, making it a more reliable choice. The decline in performance in RMS can be explained given the large firing rate dependence of the RMS noise estimate (Quiroga et al. 2004). Despite the decline in performance, RMS noise estimates are commonly used to set spike detection thresholds in MEA recordings (Ronchi et al., 2021; Sommer et al., 2022; Miyahara et al., 2023; Bertacchi et al., 2024; Vinci et al., 2025; Setsu et al., 2025; Kesdoğan, et al., 2024; Fenton et al., 2024; Hoang et al., 2025; Zhao et al., 2025). Although this method is acceptable in a burst-free context, network bursts are the main activity mode of primary culture models (Murphy et al., 1992; Wong et al., 1993; Rhoades & Gross, 1994; Kamioka et al., 1996; Canepari et al., 1997; Voigt et al., 1997; Segev et al., 2001; Harris et al., 2002; Wagenaar et al., 2006; Chiappalone et al., 2007; Chen and Dzakpasu, 2010; Colombi et al., 2013; Suresh et al., 2016).

Because of their similar approach, the RMS and MED methods have a directly comparable output, which is the number of spikes detected across channels. Both methods detect spikes on individual channels; however, they are unable to distinguish duplicated spikes that may arise from multiple channels. On the other hand, KS4 and other spike sorting methods create templates from spike waveforms across multiple channels (Pachitariu et al. 2024). Therefore, direct comparison of RMS and MED methods with sorting algorithms is difficult, requiring ground truth data. To accomplish this, we simulated data that is similar to HD-MEA recordings using LFPY and mearec python packages (Lindén et al., 2014; Buccino & Einevoll, 2021). The simulated data emulates a mature cortical culture after three weeks *in vitro*, showing spontaneous activity and network bursts as we observed in our HD-MEA recordings. Evaluation of precision on the simulated data indicated that DP-MED is more precise than KS4 when firing rate is high, but KS4 has better recall. This indicates that at our chosen parameters, KS4 is more likely to over-detect or mis-assign spikes to units during periods of intense activity. On the other hand, DP-MED, while less sensitive to firing rate than RMS, still raises its threshold during intense periods of activity and will therefore be more likely to miss spikes. Previous work has shown that KiloSort2 can detect more false positives than a multi-sorter consensus framework based on retaining spikes found by multiple sorting algorithms (Buccino et al., 2020), although that estimate is highly dependent on the shape of the action potential (Pachitariu et al., 2024). Overall, the differences between DP-MED and KS4 reflect a tradeoff between precision and recall, and both methods are comparable in performance.

Last, we tested the performance of DP-MED and KS4 on conditions similar to those that may be encountered during drug screening trials, using the HD-MEA platform. We used carbachol to pharmacologically stimulate bursting patterns and spontaneous activity through activation of nicotinic and muscarinic receptors (Tateno et al. 2005a; Tateno et al. 2005b; Hammond et al. 2013). As expected, we observed that spontaneous activity and network burst frequency increased with the addition of carbachol, regardless of the detection method used. We did not see a statistic significant difference between the number of spikes detected by DP-MED and KS4. However, there was a tendency for KS4 to detect more spikes, which is consistent with our simulated data analysis. KS4 detects more false positives and fewer false negatives, increasing the number of spikes detected. Overall, we showed that both methods are comparable when tested on simulated data and in a non-simulated biological context. Nevertheless, a critical difference is that the template generation and sorting steps make sorting algorithms like KS4 computationally heavier than DP-MED. This difference could be even more noticeable in high throughput pharmacology approaches; where the spike density increases during recordings. Here, we showed that, when relying solely on a CPU, DP-MED is 40 times faster than KS4 in processing HD-MEA recordings of ∼1000 channels over 900 seconds at 20 kHz. Furthermore, the runtime of DP-MED was a fraction of the real recording duration, making online detection possible. To our knowledge, this is the first study that proposes a fast, easy-to-implement pipeline for HD-MEA recordings that is comparable to one of the most popular open-source sorting algorithms, KS4.

In this research, we showed how commonly used RMS-based threshold detectors underestimated the number of spikes during network burst periods. We propose a new pipeline of spike detection that combines the MED detection method with a heuristic de-duplication step. This algorithm is easy to implement, simple, and fast for detection of EAP in HD-MEAs recordings. The DP-MED method is less computationally expensive than an open-source sorting algorithm such as KS4. The performance of DP-MED is comparable to KS4 in terms of performance, offering a trade-off between precision, recall, and accuracy. However, this method is forty times faster than the implementation of KS4 in HD-MEA recordings. DP-MED performed similarly in a pharmacology experiment in which the cultures were stimulated by the addition of carbachol, a broad-spectrum cholinergic agonist. Thus, this method offers a fast and accurate interpretation of HD-MEA in drug screening, pharmacology, and fundamental neuroscience applications.

## 5. Acknowledgments

This work was supported by the National Science Foundation GRANT-2133769 ‘URoL-Learning the Rules of Neuronal Learning’. We thank Lauren Gomez for helpful comments and Aditi Kulkarni for technical assistance.

## 6. Data availability

Data will be made available on request. The full pipeline is available on GitHub (https://github.com/rol-group/med-dedup-detector/tree/main)

