## Supplemental Figures for "A fast and simple algorithm for accurate spike detection in HD-MEA recordings"

**10. Supplementary figures**

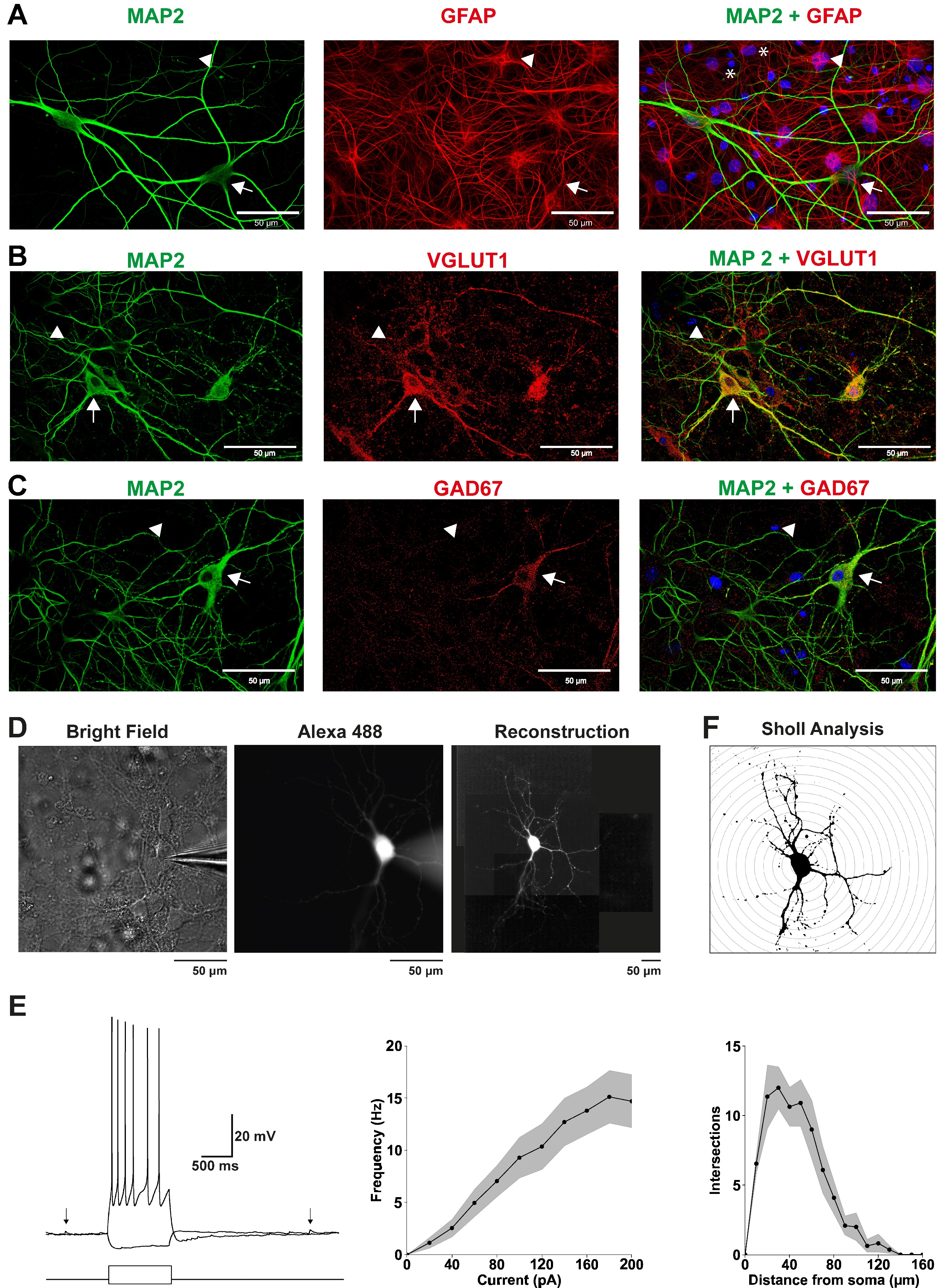

**Supplementary figure 1. Characterization of cultured primary cortical neurons**. **A.** Three weeks after seeding, cells were fixed and immuno-stained for MAP2, uniquely found in neurons; GFAP, uniquely found in astrocytes, and DAPI (shown in blue). The merged images show that some nuclei correspond to neurons (white arrows) and some nuclei correspond to astrocytes (white arrowheads). Some labeled nuclei do not show either MAP2 or GFAP staining, suggesting the presence of a different subtype of glial cells or cells that did not develop (white asterisks). **C, B**. Excitatory neurons were immuno-stained for VGLUT1 and inhibitory neurons for GAD67 (both in red). Arrows indicate overlapping staining for the markers with MAP2; arrowheads indicate DAPI stained cells that do not show overlapping staining with the neuronal antibodies. **D, E & F.** Electrophysiological and Morphological characterization of cultured cortical neurons. **D**. Left, bright field image of a neuron recorded in whole-cell mode, using Alexa Fluor 488 dye in the intracellular pipette solution (middle). Right, composite image of the labelled neuron, post recording. Cells were recorded in the current clamp configuration, using K-Gluconate internal solution containing. **E.** Left, the electrical response of the cell was tested using current steps of 20 pA, starting from –20 pA. Response of the cell to –20 and 60 pA current steps. Right, plot showing the frequency of spikes elicited by current steps of increasing amplitude. Black arrows indicate the presence of post synaptic potentials (n=22, 4 cultures). **F.** Top: Sholl analysis was used to assess neuronal branching patterns. Bottom: Plot showing the number of branch intersections with concentric rings spaced at 10 μm intervals from the cell body (n = 11 cells, 3 cultures). Scale Bar is 50 µm.

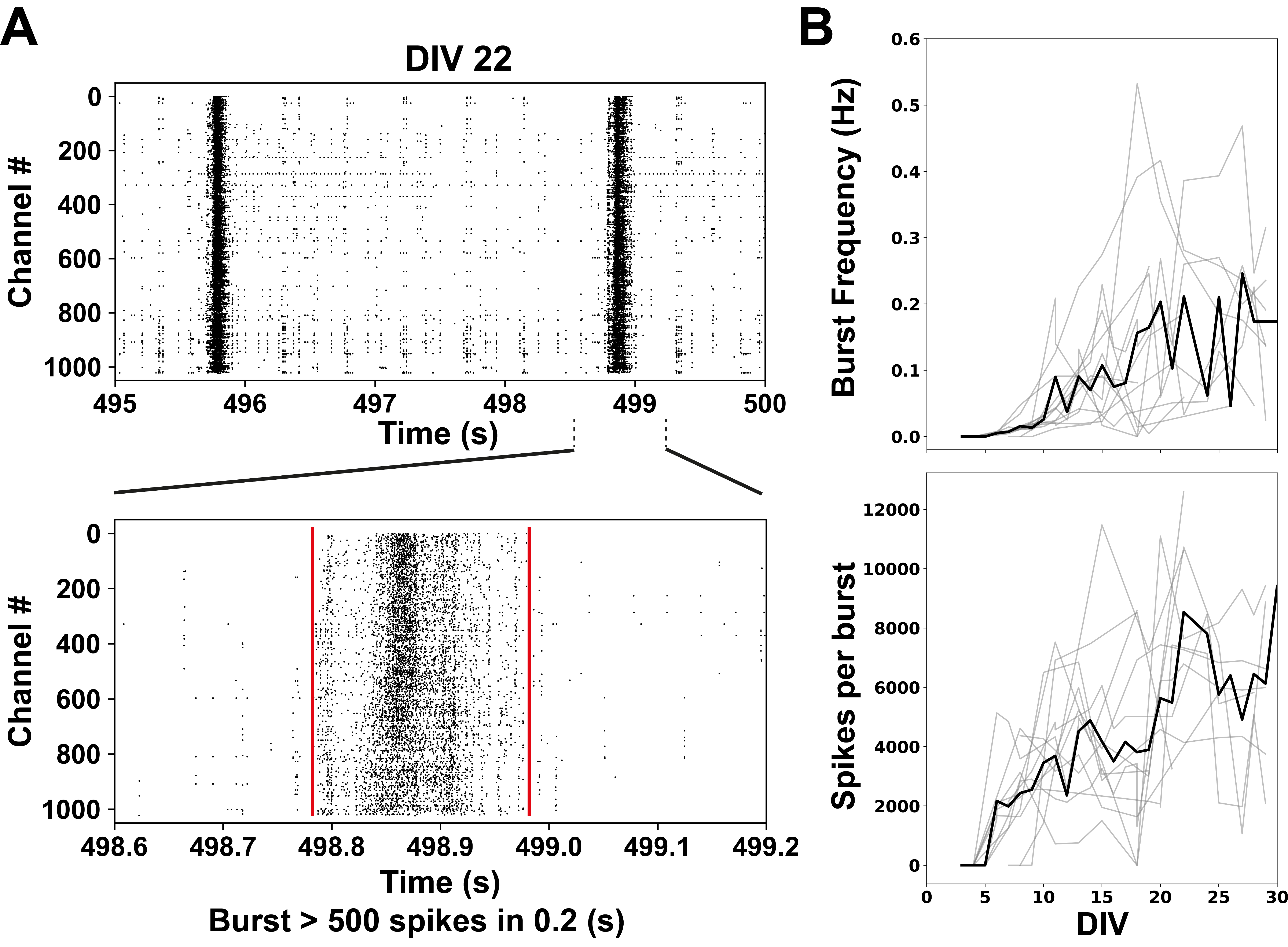

**Supplementary figure 2. Characterization of primary cortical cultures development in HD-MEAs**. **A.** Top, time window expanded from the raster plot at DIV22 shown in Figure 1B. Here we illustrate spontaneous activity and thick bands of synchronized activity that we will characterize as network bursts. Bottom, time window highlighted in the upper raster plot with a black line to show a network burst better defined in the temporal scale. Network bursts are defined as an event of 0.2 s and at least 500 spikes. **B.** Two different network burst properties evaluated across DIV. Top, burst frequency increases until ~DIV 22, where it reaches a plateau. Bottom, spikes per burst increases until ~DIV 22, where it reaches a plateau (n=11 cultures).

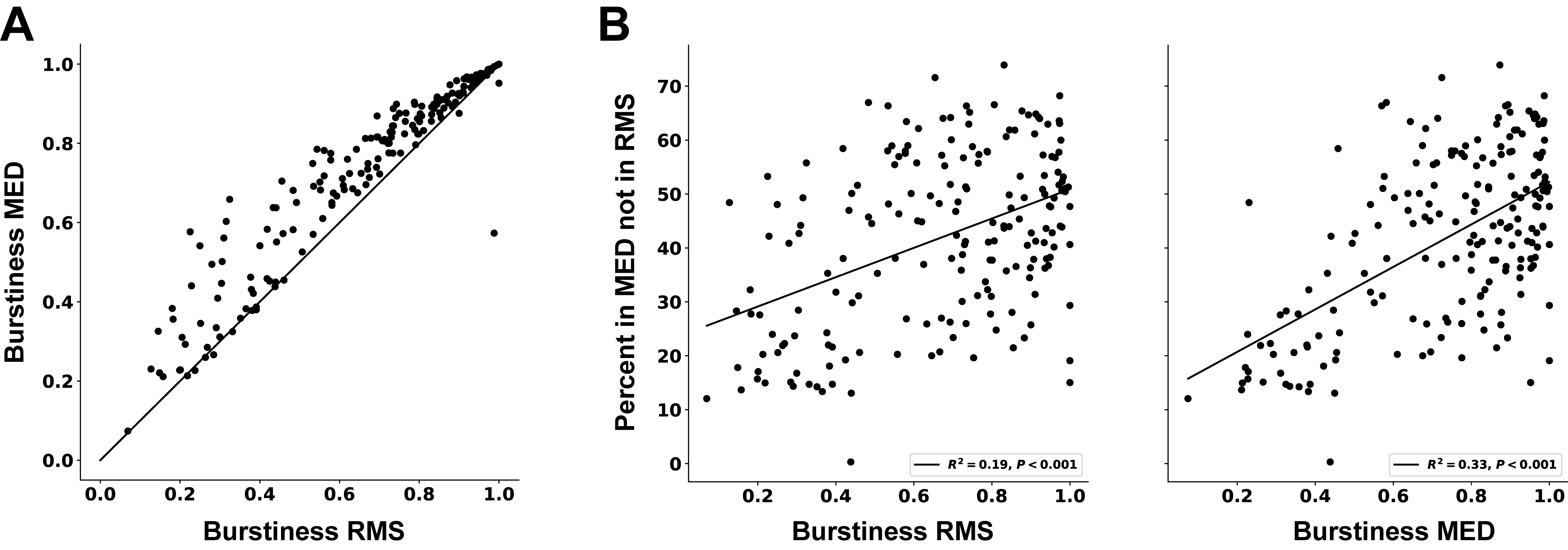

**Supplementary figure 3. Detector Similarity as a Function of Burstiness.** **A.** Burstiness. The fraction of spikes in the top 15% of 0.5s time bins is measured relative to chance and compared for each detection method. This metric is called burstiness. Each dot is a recording. The line shows unity. (N = 55 Recording Sessions). **B.** Detection Overlap and Burstiness are Correlated. The percentage of spikes unique to the MED detection is plotted against the burstiness of each recording. Left: Burstiness values computed from RMS spike trains are used. (OLS: R^2^ = 0.32; ANOVA F-test: F_1,53_ = 24.93, P = 6.8 x 10^-6^) Right: Burstiness values computed from median spike trains are used. (OLS: R^2^ = 0.29; ANOVA F-test: F_1,53_ = 21.23, P = 2.6 x 10^-5^).

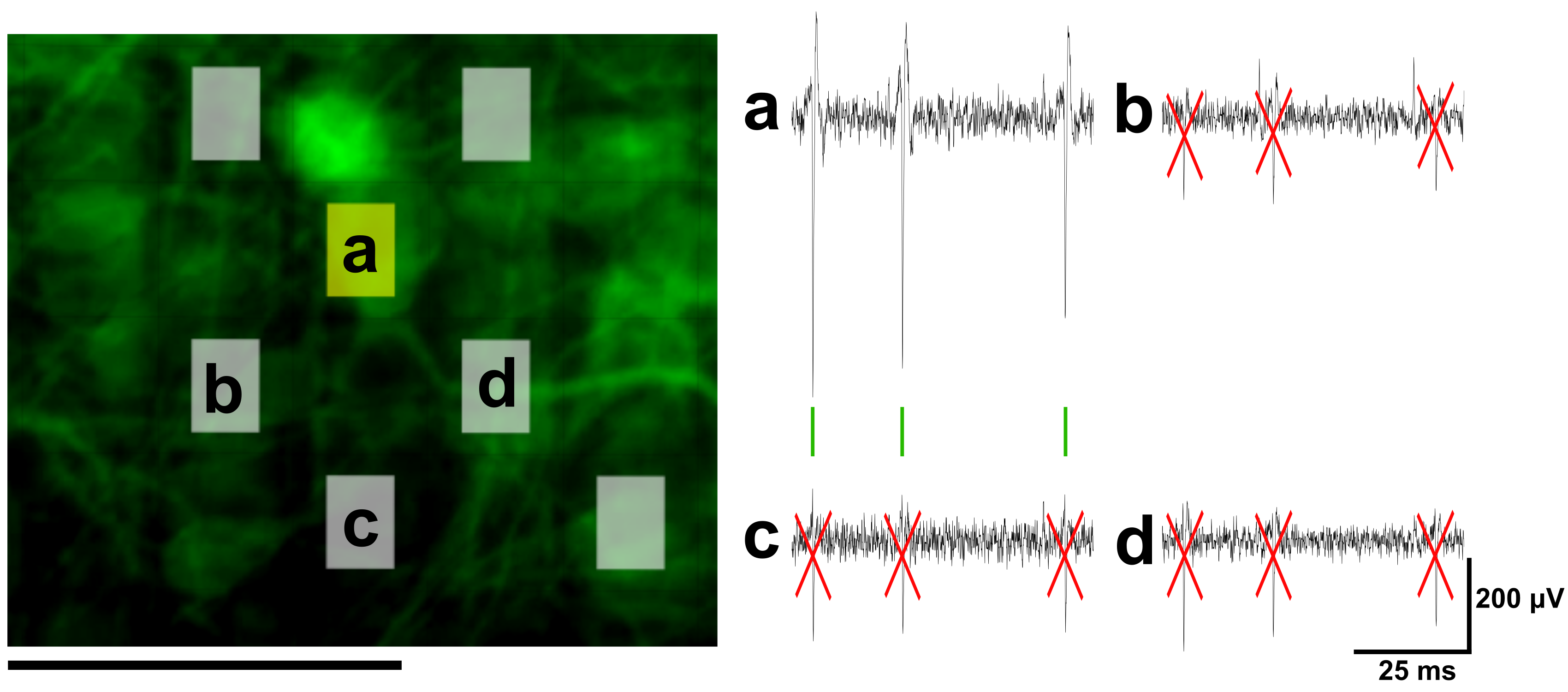

**Supplementary figure 4. Visual representation of DP-MED spike detection method.** Left: Epifluorescence image of an HD-MEA chip containing cortical neurons, fixed post-recording, immunolabeled for MAP2 (green). Superimposed on this image, we show the electrodes recorded. In yellow, we highlighted the electrode that recorded highest amplitude events from a sub-set of 4 electrodes (a-d). The yellow electrode is on top of a well-defined cell body with processes. Scale Bar is 50 μm. Right: Filtered voltage traces (300 Hz high pass) recorded by 4 electrodes (a-d). First, the spikes were identified using the MED threshold detection method. The high amplitude spikes (green) were recorded by surrounding electrodes with a smaller amplitude (b-d). De-duplication step eliminates smaller amplitude detections (illustrated as red x) and keeps the high amplitude ones (green). Scale Bar is 200 μV and 25 ms.
